# Identification of Mechanisms of Functional Signaling Between Human Hippocampus Regions

**DOI:** 10.1101/099820

**Authors:** Ruben Sanchez-Romero, Joseph D. Ramsey, Jackson C. Liang, Clark Glymour

## Abstract

**Background:** Standard BOLD connectivity analyses depend on aggregating the signals of individual voxels within regions of interest (ROIs). In certain cases, this spatial aggregation implies a loss of valuable functional and anatomical information about subsets of voxels that drive the ROI level connectivity.

**New Method:** We use the FGES algorithm, a data-driven score-based graphical search method, to identify subsets of voxels that are chiefly responsible for exchanging signals between ROIs. We apply the method to high-resolution resting state functional magnetic resonance imaging (rs-fMRI) data from medial temporal lobe regions of interest of a single healthy individual measured repeated times over a year and a half.

**Results:** The FGES algorithm recovered subsets of voxels within larger medial temporal lobe ROIs of entorhinal cortex and hippocampus subfields that show spatially consistency across different scanning sessions, and are statistically significant under tests that validate the role of these subsets as main drivers of effective connectivity between hippocampal regions of interest.

**Comparison with Existing Methods:** In contrast to standard functional connectivity methods, the FGES algorithm is robust against false positive connections produced by transitive closures of adjacencies (correlation methods) and common effect conditioning (Markov random field methods).

**Conclusions:** The FGES algorithm allows for identification of communication subsets of voxels driving the connectivity between regions of interest, recovering valuable anatomical and functional information that is lost when ROIs are aggregated. The FGES algorithm is specially suited for voxelwise connectivity research, given its short running time and scalability to big data problems.

## 1. Introduction

To improve the signal-to-noise ratios in relatively small samples yielding a few hundred timepoints of 50,000 or more cortical voxels, fMRI measurements, recorded at approximately 2x2x2 mm spatial resolution, have commonly been clustered into a few regions of interest each consisting of hundreds or thousands of such voxels (Nieto-Castanon et al., 2003; Faria et al., 2012; Wong 2014). The averaged measurements are then taken as an indication of activity—or its absence—engaged by some task or by the brain at rest, and the correlations or partial correlations of the averaged signals have been used as estimates of signaling connections between regions represented as an undirected graph (Salvador et al., 2005; Hayasaka et al., 2010; Smith et al., 2013;) Appropriate criteria for clustering of voxels are disputed, but in some cases, such as the medial temporal lobe, there are well-developed anatomical, histological and experimental grounds for distinguishing neural tissues among which there are signaling connections, as in Figure 1.

**Figure 1.**
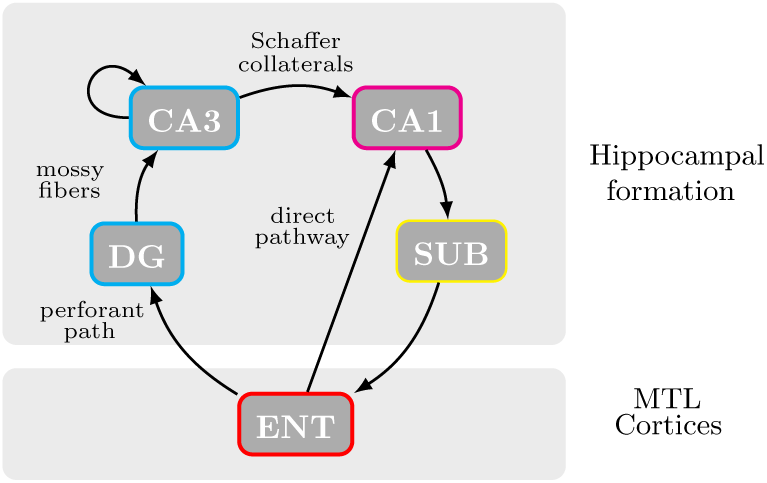
One interpretation of ground truth for the medial temporal lobe effective connectivity, including MTL cortices (entorhinal cortex (ENT)) and hippocampal formation (dentate gyrus (DG), CA3, CA1 and subiculum (SUB)).

### 1.1 Communication Subsets Model

It seems unlikely that neurons in different voxels within such anatomical regions of interest all act homogeneously in sending and receiving signals to and from neurons in voxels in other anatomical regions of interest. An alternative, more mechanical model postulates that for each pair of regions that signal one another directly--via a channel without any intermediate third region--some subsets of “communicating” voxels in the two respective regions are principally involved in sending and/or receiving. Two (or more) sets of voxels within a region that communicate respectively with two (or more) other regions should have different, but possibly overlapping, communicating subsets as suggested in Figure 2. Voxels within a region that are in none of its communication sets may serve as transmitters between them.

**Figure 2.**
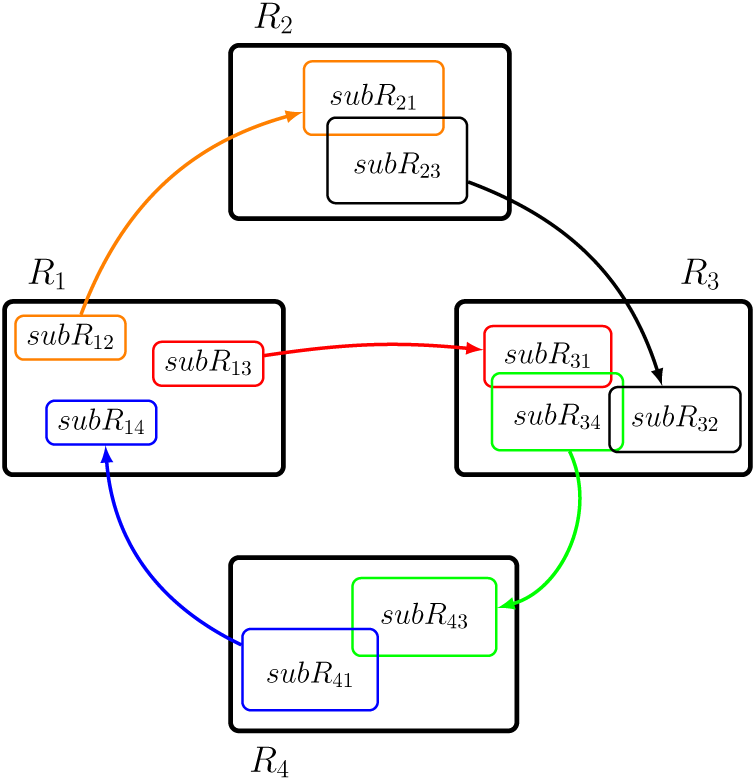
Mechanical model postulating the presence of communication subsets of voxels *subR*, within larger regions of interest *R*, driving the connectivity between regions. Communications subsets *subR* are distinct but can overlap.

This model suggests more specific hypotheses: (*H1*) for any two directly connected regions there should be respective proper communication subsets of voxels that, when averaged, remain strongly correlated when all other voxels in the two regions are removed from the data; (*H2*) comparably sized and compact alternative subsets (not including voxels in the relevant estimated communication subsets) within the respective regions should not show strong correlations of average BOLD signals, when all other voxels in the two regions are removed from the data; (*H3*) the removal from the data of the pairwise communication subsets should leave the averaged signals of the remaining voxels uncorrelated or nearly so; (*H4*) removal of comparably sized and compact alternative subsets (not including voxels in the relevant estimated communication subsets) within the respective regions should not produce very much reduced correlations of average BOLD signals; (*H5*) the subsets of voxels with which one region communicates with two (or more) others should be distinct (in the set theoretic sense) but correlated.

The recent availability of a long sequence of resting state fMRI scans of the same individual makes it possible to use appended scans over multiple scanning sessions in place of voxel averaging across individual sessions. The availability of repeated resting state scans for the same individual permits multiplying the sample size by appending scans, while reducing concerns about co-alignment of scans and false positive associations due to mixing of distributions from scans of different individuals. High-resolution voxel level measurements in turn make it possible to test these hypotheses on the medial temporal lobe using machine learning methods.

## 2. Materials and Methods

### 2.1 Data

The data used here were acquired, preprocessed, and provided to us by Russell Poldrack as part of the MyConnectome Project (myconnectome.org). We provide general information about the acquisition and preprocessing. Full details are available in Laumann et al. (2015) and Poldrack et al. (2015). MRI data were obtained repeatedly from one healthy individual over the course of 18 months. Scanning was performed in a fixed schedule, subject to availability of the participant. Scans were performed at fixed times of day; Mondays at 5 pm, and Tuesdays and Thursdays were performed at 7:30 am. Imaging was performed on a Siemens Skyra 3T MRI scanner using a 32- channel head coil. T1- and T2-weighted anatomical images were acquired using a protocol patterned after the Human Connectome Project (Van Essen et al., 2012). Anatomical data were collected on 14 sessions through 4/30/2013, with a one-year follow up collected on 11/4/2013. T1-weighted data were collected using an MP-RAGE sequence (sagittal, 256 slices, 0.8 mm isotropic resolution, TE=2.14 ms, TR=2400 ms, TI=1000 ms, flip angle = 8 degrees, PAT=2, 7:40 min scan time). T2-weighted data were collected using a T2-SPACE sequence (sagittal, 256 slices, 0.8 mm isotropic resolution, TE=565 ms, TR=3200 ms, PAT=2, 8:24 min scan time). Resting state fMRI was performed using a multi-band EPI (MBEPI) sequence (Moeller et al., 2010) (TE = 30 ms, TR=1160 ms, flip angle = 63 degrees, voxel size = 2.4 mm x 2.4 mm x 2 mm, distance factor=20%, 68 slices, oriented 30 degrees back from AC/PC, 96 x 96 matrix, 230 mm FOV, MB factor=4, 10 min scan length). Starting with session 27 (12/3/2012), the number of slices was changed to 64 because of an update to the multiband sequence that increased the minimum TR beyond 1160 ms for 68 slices. A total of 104 resting state fMRI scanning sessions were acquired; 12 were pilot sessions using a different protocol, and additional 8 were excluded based on poor signal, leaving a total of 84 usable sessions. Functional data were preprocessed including intensity normalization, motion correction, atlas transformation, distortion correction using a mean field map, and resampling to 2mm atlas space. No spatial smoothing was applied. For all sessions, the last 37 volumes were removed due to a preprocessing artifact that produced unusually high measures of the BOLD signal, leaving a total of 480 volumes per scanning session.

### 2.2 Regions of Interest

Regions of interest in the medial temporal lobe were defined manually according to procedures established by the Preston Laboratory at The University of Texas at Austin (Liang et al., 2013). ROIs were defined bilaterally for subiculum, CA1, CA32DG and entorhinal cortex. The high-resolution T2 anatomical images of the MyConnectome Project allowed a more reliable delineation of hippocampal subfields in the body, head and tail of the hippocampus; Insausti and Amaral (2004) and Duvernoy et al., (2013) were used as anatomical guidelines. Regions for the left hemisphere are illustrated in figure 3. Regions for the right hemisphere are shown in supplementary figure S1.

**Figure 3.**
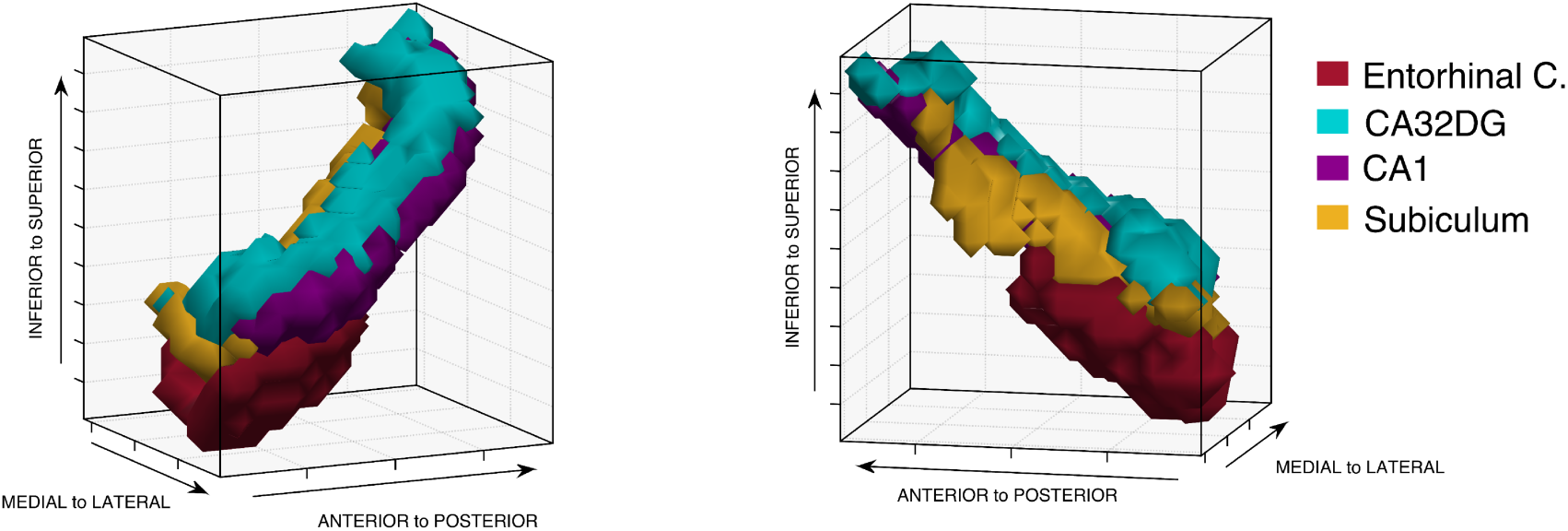
Lateral view (left) and medial view (right) of 3D rendered left hemisphere medial temporal lobe showing ROIs for entorhinal cortex and hippocampus subfields.

The dentate gyrus (DG) is hardly separable in functional MRI from CA3 and CA2, resulting in the comprehensive ROI labeled CA32DG (Zeineh et al., 2000; Ekstrom et al., 2009; Preston et al., 2010). The entorhinal cortex is also a complex structure for which several subregions have been distinguished in the literature but are not distinguished in our data (Kerr et al., 2007; Canto et al., 2012). Simulation results (Smith et al., 2011) suggest that effective connections involving such confounded regions may be difficult to identify.

### 2.3 Replication

For statistical and replication analysis, four datasets were created for each hemisphere by appending: one subset of the first five consecutive sessions; two subsets of ten consecutive sessions, the first ten and the last ten; and one subset of ten random sessions.

### 2.4 Methods for Identification of Communication Subsets

Several kinds of procedures are available to estimate voxel to voxel connections, including simple correlation, penalized inverse covariance—which is effectively penalized partial correlation controlling for all variables—vector or structural autoregression, PC, a constraint-based graphical search method (Spirtes and Glymour, 1991; Spirtes et al., 2000), and the score-based graphical search method we used here, FGES (described in Ramsey, 2015).

FGES is a fast implementation of the quasi-Bayesian GES algorithm (Meek, 1997; Chickering, 2002; Chickering and Meek, 2002; Ramsey et al., 2010; Mumford and Ramsey, 2014), with altered caching and parallelization, known to return asymptotically the correct adjacencies from i.i.d. data generated from a directed acyclic graph with a Gaussian distribution of its variables, represented as graph vertices. Two temporally close BOLD measurements of a voxel are of course not independent, but the great majority of measurements from a voxel will be widely separated from one another in time and can be regarded as independent. In simulations, FGES tolerates small deviations from Gaussian distributions and i.i.d sampling. We use only the adjacencies estimated by FGES, not the estimates of directions of influence, which are less reliable. The FGES algorithm is publicly available as part of the TETRAD causal discovery freeware suite (www.phil.cmu.edu/tetrad/), and Java code is available in the TETRAD open source project, which can be cloned at (github.com/cmu-phil/tetrad).

FGES uses the Bayes Information Criterion (BIC) to assess models (Schwarz, 1978). The score is the difference between a penalty that depends on number of parameters and sample size, and the log-likelihood of the model, with lower scores preferred. The penalty can be adjusted. The high penalty, 30, we have used in the BIC score filters out comparatively weak partial correlations. At a much lower penalty, 4, for example, little or no separation of voxel subsets is discovered by FGES. Weak partial correlations can be expected between voxels that have no direct influence on one another because statistically controlling for the BOLD signal of voxels physically intermediate on a channel between two voxels cannot fully control for propagated influences, as illustrated in Figure 4, where *X, Y* and *Z* denote neuronal physiological activity in a voxel region, *BOLD* denotes the corresponding recorded BOLD signals, *S* denotes exogenous inputs and *E* denotes measurement errors. Were *X*, *Y*, and *Z* measured precisely, *X* and *Z* would be independent controlling for *Y*, but *BOLD_X_* and *BOLD_Z_* will not be strictly independent controlling for *BOLD_Y_*.

**Figure 4.**
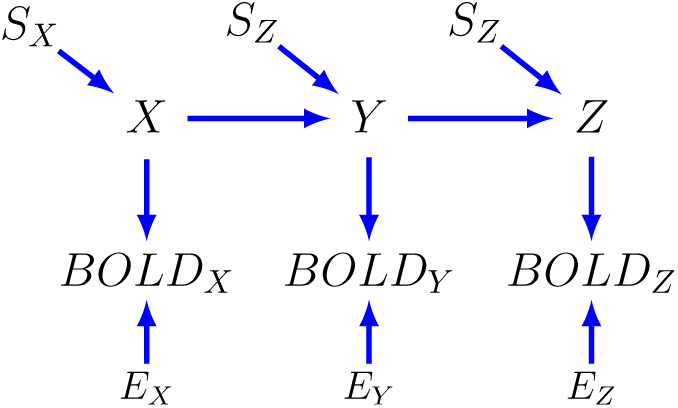
Indirectly measured causal structure relating physiological activity and BOLD recordings.

Correlation methods produce transitive closures of adjacencies rather than adjacencies. Thus in figure 4, correlation would always find an adjacency between *BOLD_X_* and *BOLD_Z_*, and were there multiple pathways between *X* and *Z*, the correlation of *BOLD_X_* with *BOLD_Z_* could be larger than the correlation of *BOLD_X_* with *BOLD_Y_* or *BOLD_Y_* with *BOLD_Z_*.

Markov random field methods, such as partial correlation and penalized inverse covariance, explicitly or implicitly condition on all variables, reducing power. These methods do not return asymptotically the adjacencies in a true graphical causal representation of the data generating process; instead, even if the variables are perfectly measured, they introduce a false adjacency between variables that are not directly connected (not adjacent) but that are both direct causes of any third variable, as in figure 5.

**Figure 5.**
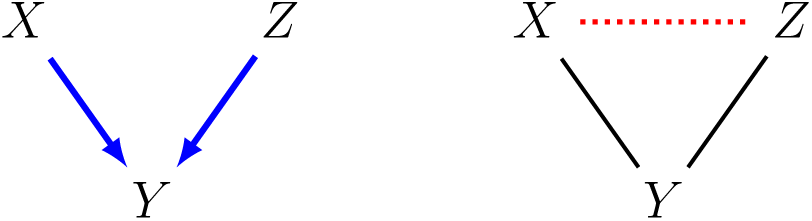
Graphical structure where variable X and variable Z are both direct causes of variable Y (left). Under this causal structure, Markov random field methods will estimate a false adjacency (red dotted line) between X and Z.

While they have worked well in simulations of fMRI data from small, sparse structures (Smith et al., 2011), in real biological and physical applications Markov random field methods have shown poor separation of regions (e.g., of regulatory versus non-regulatory genes or surfaces of different mineral composition) known to have distinct causal roles in generating the data (in the examples, respectively, phenotype and reflectance spectrum). In our data, penalized inverse covariance with a very high sparsity requirement (using the QUIC implementation (Hsieh et al., 2013)) produces dense communication subsets that are not in areas in which the hippocampal regions are in physical contact, and fails to find associations between averaged BOLD signals of the communication subsets it estimates.

Vector and structural vector autoregression methods (often called “Granger causal” methods) have shown poor adjacency detection in fMRI simulations (Smith et al., 2011). PC has generally, but not always, produced less accurate results than GES, and in current implementations is much more slower than FGES for big data problems.

In view of these considerations, we have used the FGES algorithm, which constructs a model stepwise using piecewise posterior probabilities as scores for assessing alternative steps, uses no time order information, and thanks to its altered caching and parallelization is extremely fast and can be scaled to datasets up to a million variables (Ramsey, 2015). It is of course quite possible that alternative machine learning methods, for example, methods identifying feedback relations, or functional causal methods using a time proxy as a variable would serve as well or better. We have not used them because the former are not as yet computationally feasible for problems of the size considered here, and the latter are still under development.

## 3. Results

### 3.1 FGES graphs

To obtain pairwise communication subsets for each pair of regions of interest in the hippocampus dataset, the FGES algorithm with penalty 30 was run over 570 voxels comprising the four distinct anatomical regions of interest (CA32DG, CA1, subiculum and entorhinal cortex) for the left hemisphere hippocampus (or 530 voxels for the right hemisphere hippocampus), and 4,800 (ten sessions appended) or 2,400 (five sessions appended) BOLD datapoints.

For datasets with the aforementioned dimensions, the FGES algorithm at penalty 30 in a MacBookPro 2.4 Ghz Intel Core i5, 8GB memory, returns an answer in an average of 15 seconds for 4,800 datapoints; and 7 seconds for 2,400 datapoints.

Four datasets were constructed for each hemisphere, appending the first ten sessions, the last ten sessions, ten random sessions, and the first five sessions. Four datasets and two hemispheres resulted in eight different runs of the FGES algorithm. The FGES graphs output for the left hemisphere datasets of ten sessions appended had in average, 1,466 edges (0.9% density); and 1,253 edges (0.9% density) in average for the right hemisphere datasets. For the five sessions appended datasets, the number of edges decreased to 1,051 (0.6% density) (for the left hemisphere), and 842 (0.6% density) (for the right hemisphere). The reduction in sample size affected the power of the FGES algorithm to detect functional connections. In both, left and right hemispheres, the reduction in the number of estimated edges had approximately the same magnitude.

For the three left hemisphere ten sessions appended datasets, the average mean degree of the graphs was 5.13, average maximum degree, 13.6, and minimum degree zero. For the three right hemisphere datasets, the average mean degree was 4.7, the average maximum degree 11.6 and the minimum degree zero. For the five sessions appended datasets, the mean degree was 3.68, maximum degree, 11, and minimum degree zero (for the left hemisphere); and mean degree of 3.17, maximum degree, 7, and minimum degree zero (for the right hemisphere). Degree values were highly similar across hemispheres and datasets. Complete results for number of edges and degree for the FGES algorithm results are shown in supplementary table 1.

### 3.2 Communication subsets

From the graphs output by FGES we can obtain the pairwise functional communication subsets for each desired pair of regions by selecting the voxels of a region that have *at least one* edge (direct adjacency) connecting to the other region, and respectively for the other region. Results for the communication subsets for each pair of effectively connected hippocampal regions according to figure 1, are illustrated in figure 6, for left hemisphere regions of interest and ten first sessions appended dataset. Results for the datasets formed by appending the last ten sessions, ten random sessions, and five first sessions, for left hemisphere hippocampus and right hemisphere hippocampus are shown in supplementary figures S2-S8.

**Figure 6.**
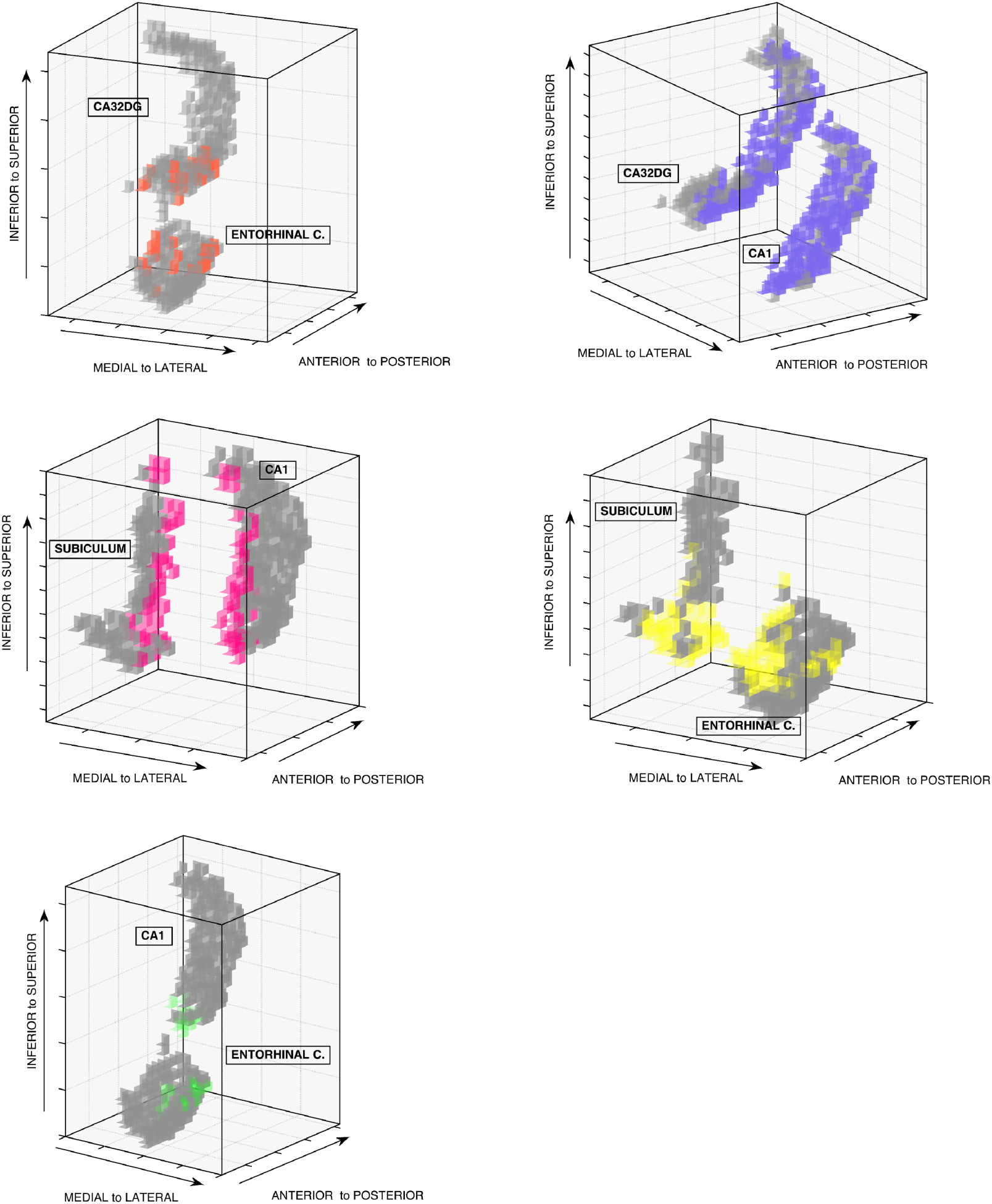
Exploded 3D views in voxel space of the communication subsets for each pair of left hemisphere hippocampal regions, estimated with voxelwise FGES algorithm for the first ten sessions appended, Entorhinal-CA32DG, CA32DG-CA1, CA1-Subiculum, Subiculum-Entorhinal, Entorhinal-CA1.

*Supplementary figures* S4 and S8 also show that for datasets of the first five sessions appended, the communication subsets are smaller. This was expected given the lower edge density and degree of the FGES graphs for these reduced sample size cases. Supplementary table 3 shows the size in voxels of the communication subsets for each pair, dataset and hemisphere. For the right hemisphere, last ten and random ten sessions appended datasets, the FGES algorithm with penalty 30 produced communication subsets with only one voxel for the pair *Entorhinal-CA1*. In these cases we can recover denser communication subsets by lowering the penalty discount of the FGES algorithm. Supplementary table 3 shows the size in voxels of the communication subsets for the pair *Entorhinal-CA1* when the FGES penalty is reduced to 12.

The communication subsets obtained using the FGES algorithm are spatially similar across the different datasets. To quantify the spatial similarity of a communication subset across different datasets we built a metric defined from 0 to 1, where larger values imply higher similarity (see supplementary table 4 for a definition of the metric and full list of results). The average of the spatial similarity metric value for all the ten communication subsets in each hemisphere across the four datasets is 0.66 (std.dev = 0.23; median=0.75; min=0; max=1; for a total of 120 similarity comparisons). Results show that bigger communication subsets are more similar across datasets, such as the subsets of the pairs *CA1-Subiculum, Subiculum-Entorhinal* and *CA32DG-CA1*, which have an average similarity metric value of 0.80. In contrast, smaller subsets are less similar, as for the pairs *Entorhinal-CA1* and *Entorhinal-CA32DG*, which have an average similarity metric value of 0.44. Of all, the *Entorhinal-CA32DG* subsets have the lowest similarity across datasets, indicating the difficulty of estimating these communication subsets with the current data.

### 3.3 Hypothesis testing for communication subsets

Hypotheses *H1* together with *H2*, and *H3* together with *H4*, of section 1.1, are reformulated here to obtain statistical tests to determine the validity of the communication subsets obtained with the FGES algorithm. These hypotheses are based on the general assumption that the communication subsets support the effective connectivity between two larger regions of interest:

*(H1) The partial correlation between two communication subsets, when all other voxels in the two regions are removed from the data, should be stronger than the partial correlation between the two complete regions.*

*(H2) The partial correlation between two alternative subsets (not including voxels in the communication subsets) when all other voxels in the two regions are removed from the data, should not be stronger than the partial correlation between the two complete regions.*

The idea behind *H1* is that the signals of the voxels contained in the communication subsets are more relevant for the effective connectivity of the regions than the signals of the rest of the voxels in the regions, and thus we expect that the average BOLD signals of the communication subsets are more correlated than when we include *all* the voxels in a region, for which the average BOLD signal will be a mixture of relevant and non-relevant voxels. Conversely, for *H2*, the connectivity of non-relevant sets of voxels is expected to be lower than the connectivity of sets that include the relevant voxels.

For each pair of regions tested, a quotient *q* is obtained by dividing the partial correlation between communication subsets over the partial correlation between the complete regions when *all* the voxels are included. If *q* > 1 there is evidence that the partial correlation between subsets is higher than the partial correlation between complete regions, as required by *H1*. Across the four datasets and the five pair of regions, the average quotient *q* is 1.44 for the left hemisphere, and 2.23 for the right hemisphere data. Full results are in supplementary table 5.

Using *H2*, we can define a permutation test to determine if an obtained value of the quotient *q* is statistically significant in a null-hypothesis permutation distribution. We build a null-hypothesis permutation distribution by first creating *N* permuted alternative communication subsets which: (i) are formed by spatially contiguous voxels; (ii) of the same size of the original communication subset obtained with FGES; (iii) do not contain voxels included in the original communication subset. We compute *N* permuted quotients *r* by dividing the partial correlation between the permuted communication subsets over the partial correlation between the complete regions when *all* the voxels are included. The *N* values of *r* form the permutation distribution of the null-hypothesis. Finally, to determine if the original quotient *q* is significantly different from the values of *r* in the null-distribution, we compute a one-tailed empirical p-value for *q*, as: *(# of r > q)/ N*. For each permutation test, we set *N*=2000.

Five pairs of regions, two hemispheres, and four different datasets imply 40 different permutation tests for *H1*: 72.5% had p-values < 0.01; 2.5% had p-values >= 0.01 and < 0.05; 5% had p-values >= 0.05 and < 0.10; and 20% have p-values >= 0.10. In particular, seven permutation tests involving the pair of regions *Entorhinal-CA32DG* were not significant, which again reflects the difficulty of detecting the communication subsets between those two regions under the current data.

All the results for the permutation tests for *H1* are in supplementary table 5.

*(H3) The partial correlation between regions for which the voxels in the communication subsets were removed, should be weaker than the partial correlation between the two complete regions.*

*(H4) The partial correlation between regions for which the voxels in alternative subsets (not including voxels in the communication subsets) were removed should not be considerably different than the partial correlation between the two complete regions.*

*H3* is motivated by the assumption that communication subsets support the effective connectivity between regions, and their removal should necessarily decrease the exchange of information. Conversely, for *H4* the removal of non-relevant subsets of voxels should not considerably affect the exchange of information between regions.

For each pair of regions tested, we obtain a quotient *q* by dividing the partial correlation between regions with the communication subsets *removed* over the partial correlation between complete regions when *all* the voxels are included. If *q* < 1 we have evidence that the partial correlation between regions with subsets removed is smaller than the partial correlation between whole regions, as required by *H3*. Across the four datasets and the five pair of regions, the average quotient *q* is 0.28 for the left hemisphere, and 0.58 for the right hemisphere. Full results are in supplementary table 6.

Using *H4*, a permutation test can be constructed for *H3* following the same steps as for *H1*, with two differences: (1) the *N* permuted quotients *r* are computed by dividing the partial correlation between the regions with the permuted communication subsets *removed* over the partial correlation between the complete regions; (2) the one-tailed empirical p-value for *q* is defined as: *(# of r < q)/ N*. For each permutation test, we set *N*=2000.

Five pairs of regions, two hemispheres, and four different datasets, imply 40 different permutation tests of *H3*: 50% had p-values < 0.01; 12.5% had p-values >= 0.01 and < 0.05; 0% had p-values >= 0.05 and < 0.10; and 37.5% have p-values >= 0.10. All the eight permutation tests for the pair *Entorhinal-CA32DG* are highly non-significant, which confirms once more the challenge of estimating the corresponding communication subsets. Three out of eight permutation tests for the pair *Entorhinal-CA1* are also highly non-significant, due to the fact mentioned in section 3.2, that the corresponding communication subsets for this pair are of size one. All the results for the permutation tests for *H3* are in supplementary table 6.

## 4. Discussion

Our results show that the communication subsets estimated with the FGES algorithm are in reasonable accord with the five hypotheses advanced in section 1.1, except for entorhinal cortex connections and especially *Entorhinal-CA32DG* connections. These exceptions are unsurprising given the inability of fMRI recordings to distinguish the dentate gyrus (DG) from CA2 and CA3, and the complexity of the connectivity of the entorhinal cortex structure. The voxel signals were not spatially smoothed during preprocessing, which should mitigate concerns that spatial proximity is principally responsible for the associations of voxels between two estimated communication subsets. The method we have illustrated could be applied to other regions of interest, but both our results and the Smith et al. (2011) simulations offer cautions that those regions must be carefully selected with due regard to what regions imaging technology can and cannot separate.

The estimated separation of communication subsets from others is not precise, because the estimates will vary with the penalization used in the search algorithm, just as conventional correlation and partial correlation results vary with threshold and significance level chosen. With repeated estimates at multiple penalties, an ordering of all voxels in any two connected regions of interest could be obtained, giving, as a function of decreasing penalty, for any voxel in one region the (average) number of voxels in another region to which it is adjacent, providing a qualitative probability ordering of voxels in the two regions. We have limited ourselves, however, to showing that with adequate sample sizes, voxelwise effective connections can be estimated and used to explore functional substructure in well-established regions of interest.

## Acknowledgments

Research for this paper was supported by a grant from the James S. McDonnell Foundation and by the National Institute of Health under Award Number U54HG008540 and Award Number 1R01LM012087-01. The content is solely the responsibility of the authors and does not necessarily represent the official views of the National Institutes of Health. We thank Russell Poldrack for sharing of the medial temporal lobe data and helpful advice.

